# Functional insights from biophysical study of TREM2 interactions with ApoE and Aβ_1-42_

**DOI:** 10.1101/2020.02.24.963264

**Authors:** Daniel L. Kober, Melissa D. Stuchell-Brereton, Colin E. Kluender, Hunter B. Dean, Michael R. Strickland, Deborah F. Steinberg, Samantha S. Nelson, Berevan Baban, David M. Holtzman, Carl Frieden, Jennifer Alexander-Brett, Erik D. Roberson, Yuhua Song, Tom J. Brett

## Abstract

**INTRODUCTION:** TREM2 is an innate immune receptor expressed on myeloid cells including microglia in the brain. How TREM2 engages different ligands remains poorly understood.

**METHODS:** We used comprehensive BLI analysis to investigate the TREM2 interactions with ApoE and monomeric amyloid beta (mAβ42).

**RESULTS:** TREM2 binding did not depend on ApoE lipidation, and there were only slight differences in affinity observed between ApoE isoforms (E4 > E3 > E2). Surprisingly, disease-linked TREM2 variants within a “basic patch” minimally impact ApoE binding. Instead, TREM2 has a unique hydrophobic surface that can bind to ApoE. This direct engagement requires the hinge region of ApoE. TREM2 directly binds mAβ42 and can potently inhibit Aβ42 polymerization, suggesting a potential mechanism for soluble TREM2 (sTREM2) in preventing AD pathogenesis.

**DISCUSSION:** These findings demonstrate that TREM2 has at least two separate surfaces to engage ligands and uncovers a potential function for sTREM2 in directly inhibiting Aβ polymerization.

## 1. Introduction

The discovery of rare, heterozygous point variants in triggering receptor expressed on myeloid cells-2 (TREM2) that significantly increase risk for the development of late onset Alzheimer’s disease (LOAD) [1, 2] has highlighted the need to comprehensively and mechanistically understand TREM2 function in the central nervous system (CNS). TREM2 is an innate immunomodulatory receptor that associates with and signals through the adaptor protein DAP12 and, in the brain, is almost exclusively expressed on microglia [3]. AD mouse models and microglia culture studies indicate that TREM2 engages extracellular ligands to transduce signals that regulate microglial functions such as inflammatory cytokine production, migration, proliferation, phagocytosis, survival, and compaction of Aβ plaques [4, 5]. In addition to the receptor form, TREM2 can also be proteolytically released [6, 7] or alternatively transcribed [8] as a soluble version (sTREM2), which is detectable in human CSF [9]. Little is known regarding the *in vivo* function of sTREM2; however, it is generally associated with beneficial phenotypes. For example, elevated levels of sTREM2 are associated with cognitive reserve in humans [10], and increasing sTREM2 in a mouse model of AD ameliorates disease pathologies, including reducing amyloid plaque load and improving functional memory [11]. Aside from modulating the function of receptor TREM2, sTREM2 could have independent functions as a cytokine-like signaling molecule [12, 13] or function in some other capacities. Altogether, these observations suggest TREM2 plays a complex and multifaceted role in CNS health that requires detailed and molecular-level understanding in order to develop potential AD treatments targeting its beneficial functions.

Structural and biophysical studies indicate that TREM2 AD risk variants (e.g. R47H, R62H [14], etc.) do not grossly impact protein structure or stability but instead likely disrupt interactions with important ligands [15, 16]. Several proteins and biological compounds have been demonstrated to bind TREM2 [17], and AD-relevant ligands such as apoptotic cells [18], phospholipids [18, 19], low- and high-density lipoprotein [20, 21], apolipoprotein E (ApoE) [22], and oligomeric Aβ [23–25] have been confirmed to induce TREM2-mediated signaling or phagocytosis. Given the multiple ligands mediating the various functions associated with TREM2, it is important to understand how TREM2 engages each of these ligands at the molecular level.

Among the many potential TREM2 ligands, ApoE and Aβ are the most provocative because of their strong associations with AD; a single copy of the ApoE4 isoform is linked to a 3-4-fold increase in LOAD risk and Aβ plaque formation is central in AD pathogenesis. Previous studies using a variety of both non-quantitative and quantitative techniques have produced conflicting results regarding the determinants of TREM2-ApoE interactions. For example, some studies have reported that lipid loading of ApoE is required for appreciable TREM2 binding [21], and some reports show that TREM2 AD risk variants (e.g. R47H) ablate or diminish interaction with ApoE [21, 26, 27], while others report the AD risk variant has no impact on lipidated ApoE binding [23]. In addition, while some studies have characterized the signaling interaction between TREM2 and oligomeric Aβ42 (oAβ42) [23–25], none have investigated any potential functional ramifications of binding to monomeric Aβ42 (mAβ42).

In this study, we undertook a comprehensive biophysical investigation of TREM2 interactions with ApoE and mAβ42 in order to conclusively characterize these interactions, map the key required residues, and elucidate new functions. We find that TREM2 binds almost equally well to unloaded or lipidated ApoE, and we identify the major hinge region of ApoE as important for TREM2 binding. We further find that the major TREM2 AD risk variants do not grossly disrupt binding to ApoE and instead map the interaction to a hydrophobic surface on TREM2 previously implicated in binding phospholipids. We also find that TREM2 binds mAβ42 and that this interaction inhibits polymerization of Aβ42, implying that one potential function of sTREM2 is to disrupt Aβ42 polymerization and plaque formation.

## 2. Results

### 2.1 TREM2 directly binds non-lipidated ApoE isoforms

We first assessed whether TREM2 or TREML2 could directly bind non-lipidated ApoE. Both TREM2 and TREML2 [28, 29] have been identified as AD risk modifiers by GWAS and are reported to have opposing functions in microglia [30]. While the endogenous TREML2 ligand is unknown, TREML2, similar to TREM2, is reported to bind anionic phospholipids and apoptotic cells [31]. However TREML2 does not bind cell-surface proteoglycans as robustly as TREM2 [15], despite also having a basic pI (hTREM2 pI = 8.5, hTREML2 pI = 9.6) and, likely, a similar structure. Thus, we chose to utilize TREML2 as a suitable reference both to evaluate the specificity of TREM2 interactions and to investigate its potential functions.

Streptavidin (SA) biosensors were used to immobilize precisely biotinylated TREMs and then test for direct binding to purified recombinant ApoE (**Fig. 1.A**). We found that both human and mouse TREM2 could bind all three isoforms of human ApoE (**Fig. 1.B-H**) (TREM2 Ig and ApoE are 74.5% and 72.5% identical between human and mouse, respectively). Kinetic fits are complicated by the fact that ApoE largely exists as a tetramer at micromolar concentrations [32]. Therefore, we determined affinities using the equilibrium responses and steady-state analysis. Human TREM2 bound ApoE4 with the highest affinity (281 nM), followed by ApoE3 (440 nM), and ApoE2 with a two-fold lower affinity (590 nM). Mouse TREM2 followed the same trends, albeit with slightly lower affinities. In contrast, human TREML2 displayed very little binding to ApoE, which could not be saturated at the concentrations tested. Thus, we observed dosedependent binding of TREM2, but not TREML2, to non-lipidated ApoE using the biolayer interferometry (BLI) system, demonstrating that TREM2 does not require ApoE to be lipidated in order to achieve binding and suggesting specific protein determinants mediate TREM2/ApoE interactions.

**Fig. 1.**
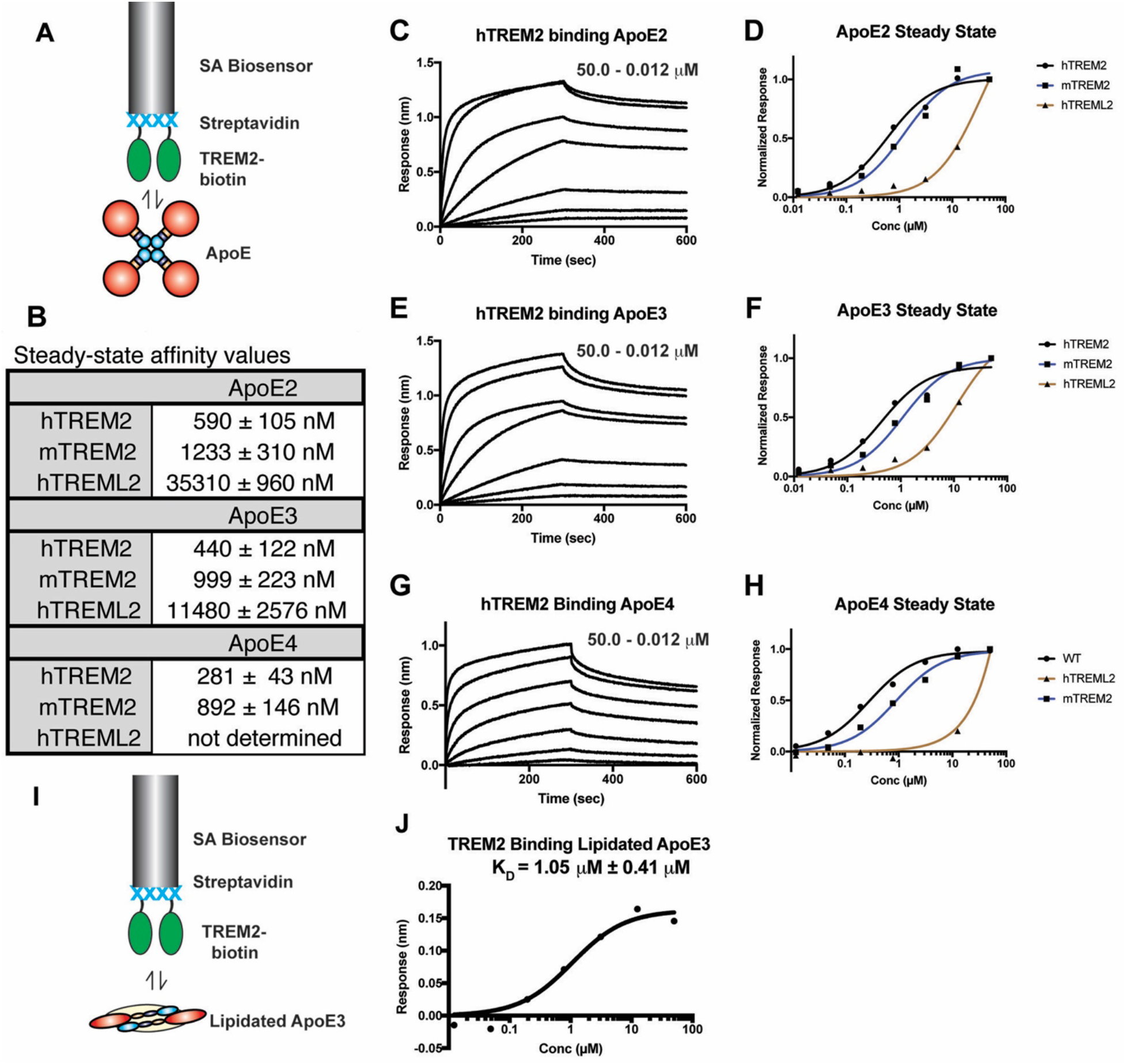
TREM2 directly binds both non-lipidated and lipidated ApoE comparably. **A)** Schematic for TREM2-ApoE BLI experiments. Biotinylated TREM2 is immobilized on a streptavidin (SA) biosensor and used to probe purified ApoE. For the experiments in C-H, the ApoE concentration range was 0.082 - 60.0 μM. **B)** Steady-state affinity values derived from the experiments shown in C-H. Affinities determined by curve fitting in GraphPad Prism. **C)** hTREM2 binding ApoE2. hTREM2 immobilized on SA biosensors were dipped in wells of ApoE2 for 300 seconds and removed and placed in buffer for 300 sec for dissociation. **D)** Steady-state responses of hTREM2, mTREM2, and hTREML2 to hApoE2. **E)** hTREM2 binding ApoE3 as in (C). **F)** Steady-state responses of hTREM2, mTREM2, and hTREML2 to hApoE3. **G)** hTREM2 binding ApoE4 as in (C). **H)** Steady-state responses of hTREM2, mTREM2, and hTREML2 to hApoE4**. I)** Schematic for lipidated-ApoE-TREM2 BLI experiments. **J)** Steady-state binding curve and affinity values of TREM2 binding lipidated ApoE3 (lipidated ApoE3 concentration range = 0.082 - 60.0 μM).

### 2.2 TREM2 binds lipidated ApoE similar to non-lipidated ApoE

Although ApoE lipidation was not required for TREM2 binding, we next tested whether lipidation might enhance the interaction. Purified ApoE3 was lipidated with DPPC *in vitro* and then re-purified by SEC for BLI experiments (**Fig. S1**). We found similar dose-dependence and overall responses with both DPPC-lipidated and non-lipidated ApoE3 (**Fig. 1.I,J**). Lipidation appeared to slightly reduce the affinity to TREM2, consistent with DPPC-ApoE particles composed of monomeric ApoE [33] compared to the non-lipidated form that is tetrameric at μM concentrations [34]. In addition, lipid composition of ApoE lipoparticles did not appear to drastically influence TREM2 binding, as astrocyte-derived ApoE4 lipoparticles bound similarly (**Fig. S2).** In summary, we observed high nanomolar to low micromolar affinity of TREM2 for non-lipidated and DPPC-lipidated ApoE3, respectively. Thus, for the remainder of our binding studies, we utilized non-lipidated ApoE.

### 2.3 TREM2 AD variants subtly alter ApoE binding while mutations to the hydrophobic patch of TREM2 significantly inhibit binding

We next assessed whether TREM2 AD risk variants alter binding to ApoE. Previous studies have not been in agreement, with some suggesting that the TREM2 R47H variant largely disrupts ApoE binding [21, 26, 27], while another showed that R47H and R62H variants retain ApoE binding [23]. We purified and biotinylated the major AD risk variants R47H and R62H as well as the T96K variant, which expands a functional “basic patch” on TREM2 that contains R47 and R62 (**Fig. 2.A, B**) and increases binding to cell-surface proteoglycans [15]. We found that R47H, R62H, and T96K only modestly reduced binding to ApoE4 (**Fig. 2.C,D**), with similar results also obtained for ApoE2 and ApoE3 (**Fig. S3**), suggesting that the basic patch on TREM2 does not largely mediate interaction with ApoE.

**Fig. 2.**
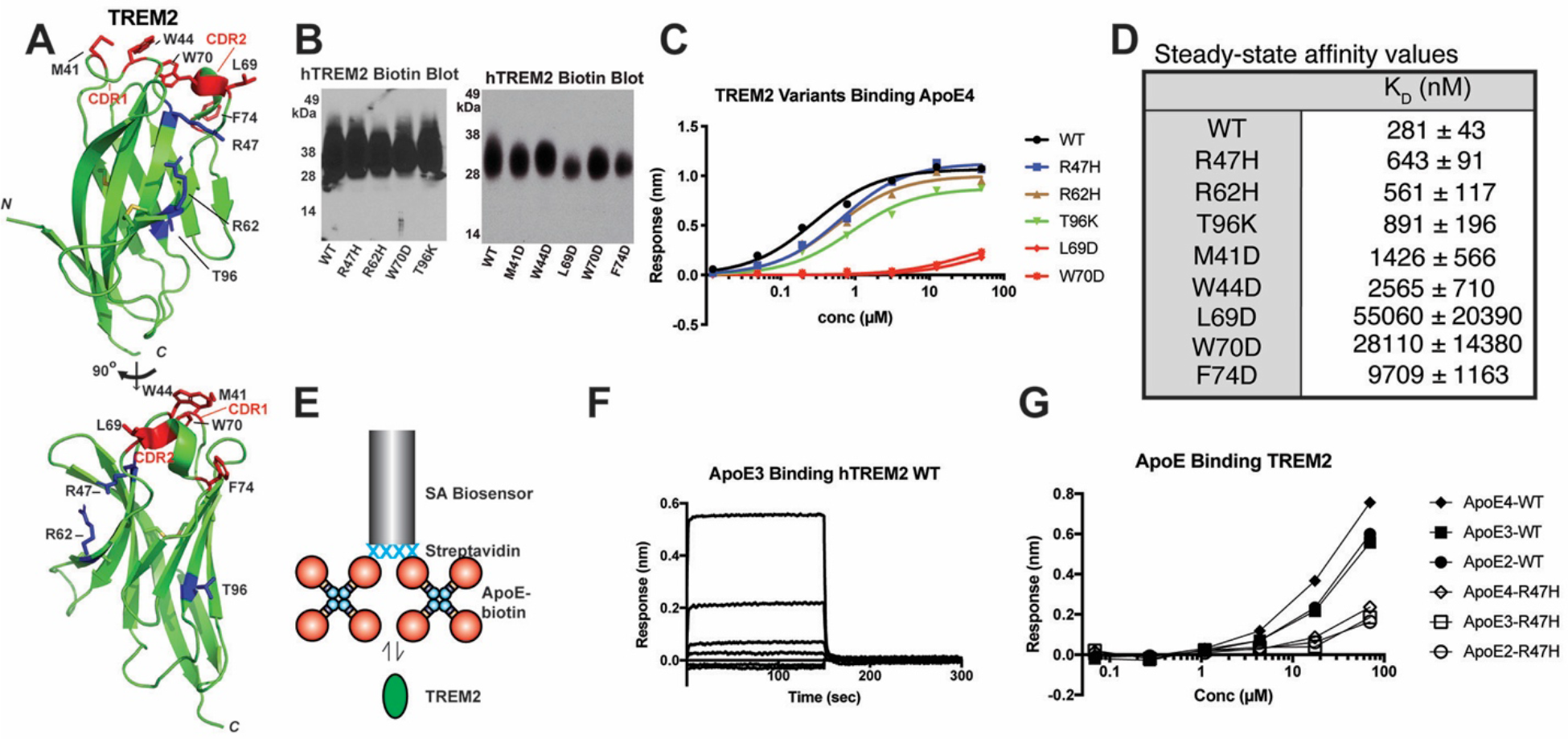
A hydrophobic patch on the distal end of TREM2 engages ApoE proteins. **A)** Crystal structure of human TREM2 (PDB 5ELI) [15] showing the positions of AD risk variants (blue) and residues in the hydrophobic patch (red) targeted in these experiments. **B)** Streptavidin-HRP blot showing roughly equivalent expression, glycosylation, and biotinylation of WT and variant TREM2 proteins. **C)** Steady-state binding of ApoE4 to WT and TREM2 variants immobilized on SA biosensors. **D)** Affinity values derived from fits of the steady state responses. **E)** Schematic for BLI experiments probing biotinylated ApoE binding to purified TREM2 proteins. **F)** Binding curves of WT TREM2 binding immobilized ApoE3. The concentration range for TREM2 was 0.068 - 70 μM. **G)** Steady-state responses of WT and R47H TREM2 binding to immobilized, biotinylated ApoE proteins.

Non-lipidated ApoE is a concentration-dependent mixture of monomers, dimers, and tetramers, with tetramers predominant at concentrations > 1 μM [34], so we wanted to address whether avidity effects from ApoE engaging multiple immobilized TREM2 molecules could potentially mask smaller changes in affinity from the R47H variant. Hence, we carried out further experiments using the opposite orientation, i.e. with ApoE immobilized and TREM2 in solution. To facilitate this, we cloned and expressed ApoE2, ApoE3, and ApoE4 with N-terminal BirA biotinylation tags and used them to probe binding to purified WT and R47H TREM2 in solution (**Fig. 2.E**). In this orientation, the binding affinity was noticeably reduced, and the binding kinetics displayed very rapid association/dissociation rates (**Fig. 2.F**). There was a noticeably decreased (but not completely ablated) response with the TREM2 R47H variant, suggesting that this mutation does slightly impact binding to ApoE (**Fig. 2.G**). However, the decrease was still relatively modest, suggesting that the basic patch in TREM2 is not the primary mediator of ApoE binding.

We next sought to identify residues on TREM2 that might directly mediate interaction with ApoE. We decided to target a hydrophobic patch that we previously identified on TREM2 [15], which is located distal to the stalk and transmembrane region (**Fig. 2.A**). Part of this region comprises a phospholipid binding surface identified in a recent co-crystal structure [16], and computational analyses have suggested that the stability of this binding surface may be disrupted by variants associated with neurodegenerative diseases, including AD risk variants [35]. Remarkably, we found that point mutations to this surface (M41D, W44D, L69D, W70D, and F74D) decreased binding affinity for ApoE4 by 5-200-fold (**Fig. 2.A-D, Fig. S4**). This finding indicates that the hydrophobic patch on TREM2 is a crucial surface mediating protein-protein interactions with ApoE. Altogether, our results demonstrate that TREM2 AD risk variants only slightly diminish binding to ApoE, while point mutations to the hydrophobic patch greatly diminish binding, strongly suggesting that this surface on TREM2 directly engages ApoE.

### 2.4 Prediction of TREM2-ApoE binding regions using a computational approach

We analyzed the sequence of human TREM2 isoform 1 using a neural-network based approach [36] and identified 19 residues most likely to be involved in protein-protein interactions (**Fig. 3.A**). Remarkably, the majority of these residues—including L69 and W70, two of the residues that strongly inhibit ApoE4 binding when mutated (**Fig. 2.C-D**)—clustered in the apical-most portion of the TREM2 ectodomain and, in particular, into three loops that compose the hydrophobic patch (CDR1: 40-47; CDR2: 67-78; and 115-120) (**Fig. 3.A**). This analysis suggested that CDR1 and CDR2 regions in TREM2 are involved in binding with ApoE.

**Fig. 3.**
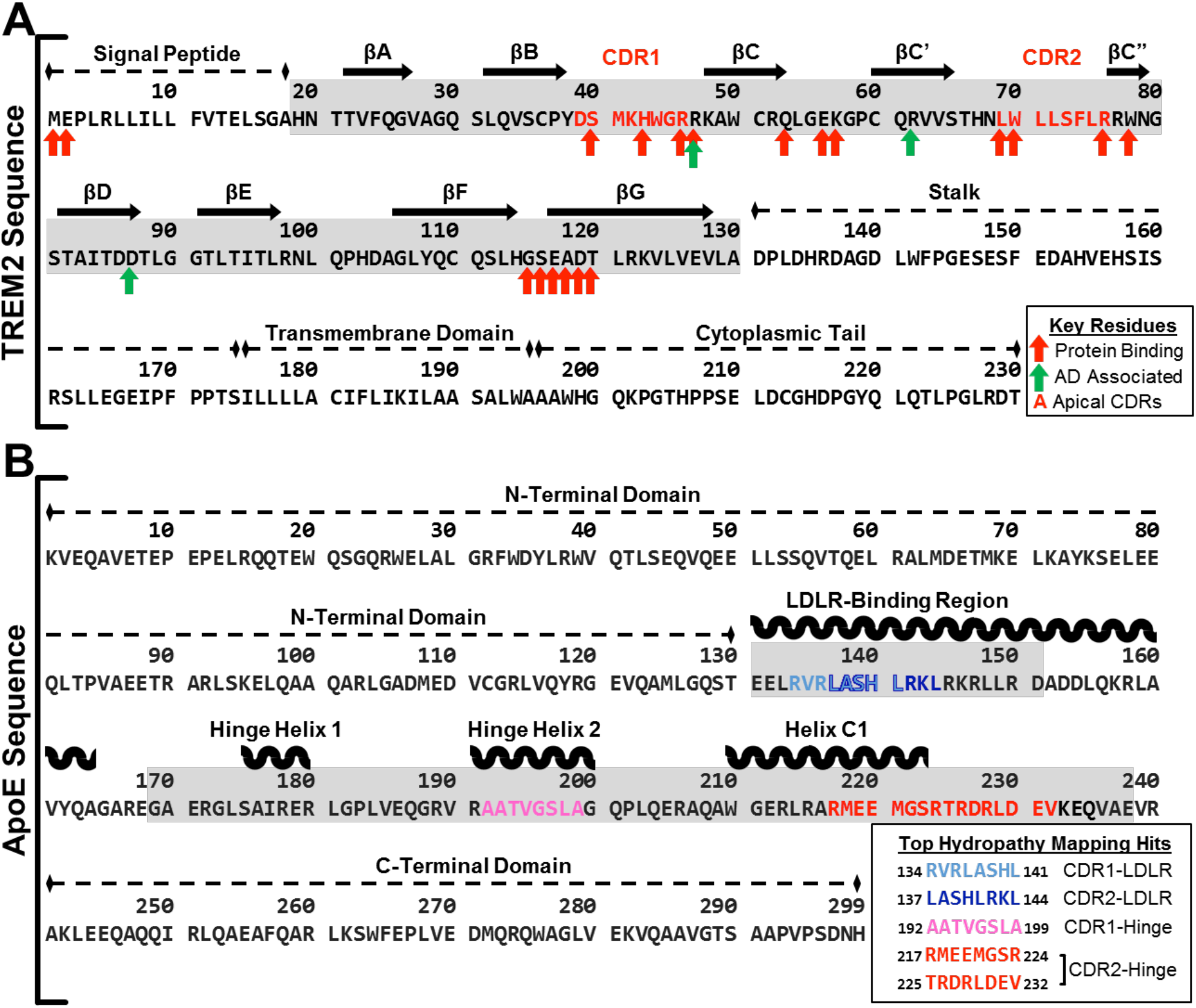
Computational analyses predict interactions between the distal hydrophobic patch on TREM2 and the hinge region of ApoE. Diagram of computationally predicted key residues and sites for protein binding on both (A) TREM2 sequence and (B) ApoE sequences. **A)** The complete sequence of isoform 1 of human TREM2 is shown with arrows to denote residues that are predicted to play a role in protein binding or disease. Red arrows denote residues predicted to act in protein-protein binding by PredictProtein software, and green arrows denote residues known to increase a carrier’s risk of developing Alzheimer’s disease when mutated. The extracellular ligand-binding domain is highlighted in gray. The sequences shown in red text represent the apical most residues of CDR1 and CDR2 used as the peptide targets for hydropathy mapping with ApoE. **B)** The complete sequence of mature human ApoE is shown with the top hydropathy mapping hits of potential binding sites for the TREM2 CDRs (colored). The two regions on ApoE where hits most heavily clustered (LDLR binding region; Hinge region) are highlighted in gray.

Since this computational approach recapitulated our experimental findings, we then used similar methods to predict the regions of ApoE which may contribute to binding this hydrophobic patch on TREM2. We used a sequence-based hydropathy mapping approach independent of ApoE conformation or lipidation state, scanning the apical-most residues of the two larger loops of the TREM2 hydrophobic surface (CDR1 and CDR2) across the sequence of ApoE separately to identify regions in ApoE that were likely to present complementary hydrophilicity profiles with the CDR1 and CDR2 loops in TREM2. Scanning in both the forward and reverse orientations of CDR1 and CDR2 in TREM2, we identified sequences in ApoE with greater than 60% hydropathic complementarity to the scanning CDR1 or CDR2 sequence that also had a palindromic partner with less than two residues difference (**Table S1**). This analysis identified the major hinge region of ApoE (192-238) that separates the N- and C-terminal domains as the region with the most consistent clustering of potential CDR-complementary sequences, with eleven sites that have more than 60% complementary hydrophilicity to both forward and reverse sequences of CDR2 and one such site for CDR1. Previously, a peptide corresponding to the receptor-binding motif in ApoE (residues 130-149) was shown to partially compete with fulllength ApoE for TREM2 binding in ELISA [22], suggesting that this region directly engages TREM2. In contrast to the hinge region, this receptor-binding motif of ApoE was found to contain only three sites that met the criteria of more than 60% complementary hydrophilicity for CDR2 and one more for CDR1 (**Table S1**). Of the complementary sequences found in the hinge region and LDLR-binding region, the three best sites (two in the hinge region and one in the LDLR-binding region) showed at least 87.5% hydropathic complementarity in both directions (**Fig. 3.B**), suggesting that the hinge region of ApoE is most likely to be involved in binding TREM2.

### 2.5 The major hinge region of ApoE contributes to TREM2 binding

In order to experimentally determine which regions of ApoE contribute to binding TREM2, we designed, expressed, and purified successive ApoE3 truncation constructs: 1) full-length ApoE3 (a.a. 1-299); 2) C-term deletion (1-238); 3) C-term and partial hinge deletion (1-191); or 4) C-term and complete hinge deletion (1-169) (**Fig. 4.A**). All the ApoE3 constructs have similar predicted pI values. All truncated proteins were purified to >95% purity by SDS-PAGE (**Fig. S5.A**) and eluted as apparent monomers by SEC (**Fig. S5.B,C**). Removal of the C-term domain greatly diminished the magnitude of the BLI binding signal (**Fig. 4.C**), largely reflecting the loss of oligomerization and, to a smaller extent, avidity, since full-length ApoE3 is a tetramer while ApoE3 1-238 is a monomer (**Fig. S2.B** and [32, 37]). Because ApoE3 1-238 is a monomer, we were able to accurately model the kinetic data 1:1, revealing a K_D_ = 696 ± 27 nM (compared to 440 nM for full-length ApoE3) (**Fig. 4.C**). In contrast, truncation into the major hinge region of ApoE3 dramatically reduced binding to barely measurable levels (**Fig 4.D.E**). These data validate our computational predictions that the major hinge region of ApoE3 (mainly residues 192-238) contributes significantly to TREM2 binding.

**Fig. 4.**
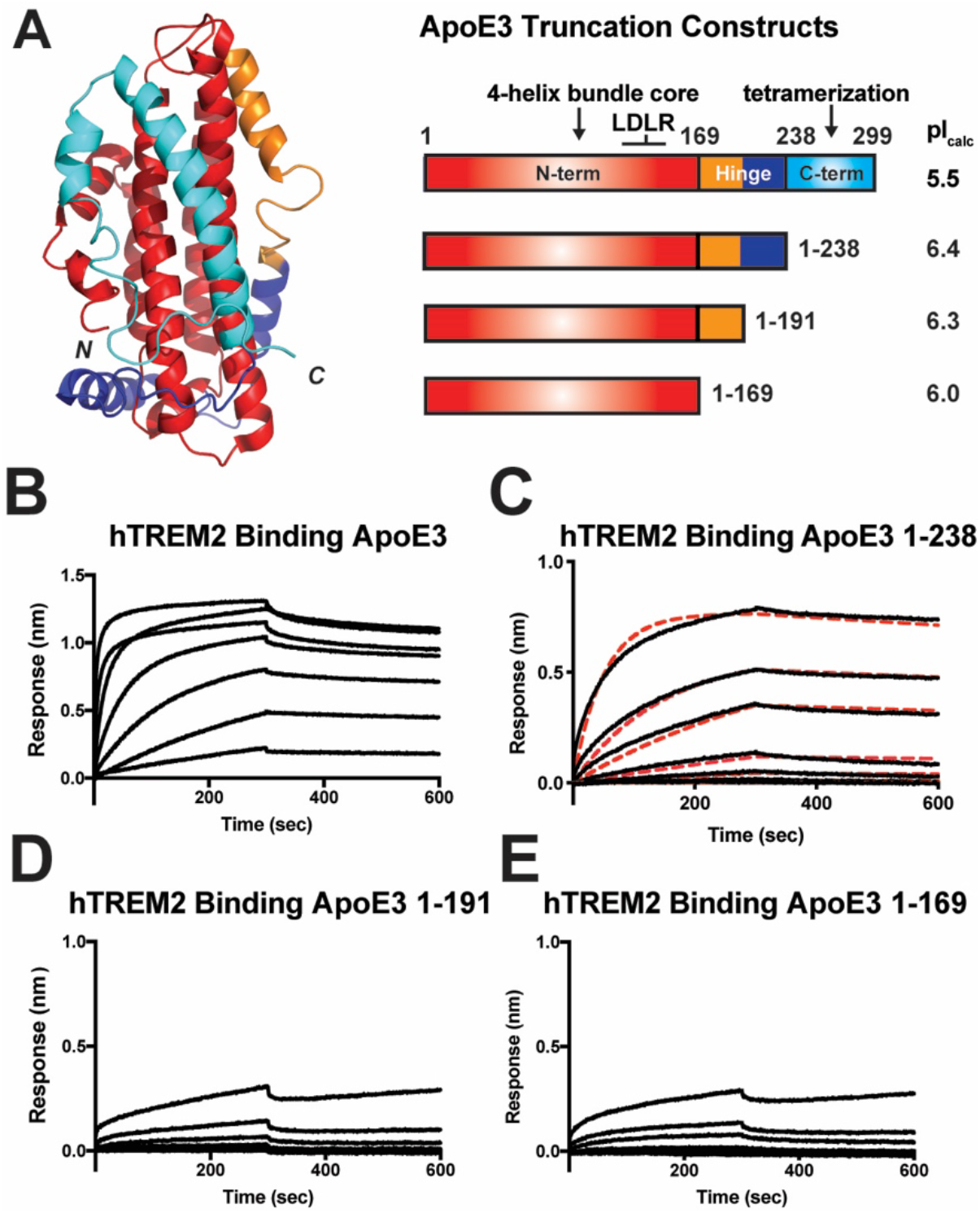
The ApoE hinge domain contributes to TREM2 binding. **A)** Schematic for ApoE truncation constructs based on structural boundaries. The NMR structure of full-length monomeric ApoE3 (PDB 2L7B) [53] is shown on the left colored according to domain boundaries in constructs shown on the right. Calculated pI values for each construct are listed on the right. (For the truncation constructs, the calculated pI values include the 8xHis purification tag). **B-E)** BLI sensograms from the experiments shown in (B). Concentration range for TREM2 in these experiments was 0.082 - 60.0 μM. Data representative of two independent experiments. **C)** BLI sensograms (black) shown with 1:1 kinetic model fits (red dash) giving a K_D_ = 696 ± 27 nM for TREM2 - ApoE3(1-238).

### 2.6 TREM2 directly binds monomeric amyloid beta_1-42_ and inhibits self-polymerization

It was recently reported that oligomeric Aβ_1-42_ (oAβ42) directly binds TREM2 and elicits TREM2-dependent signaling in microglia [23–25]. However, binding to monomeric Aβ42 (mAβ42) was not rigorously examined. In order to investigate whether TREM2 could bind mAβ42, we expressed and purified mAβ42 as a GST-fusion protein, immobilized it or free GST at equivalent levels on anti-GST biosensors, and then assessed binding to purified, monomeric TREM2 (**Fig. 5.A, Fig. S6**). In this system, TREM2 bound Aβ42 with low micromolar affinity (**Fig. 5.B,C**). Given this observation, we next investigated whether TREM2 could alter Aβ42 polymerization using an established fluorescence quenching assay [38]. Remarkably, we found that TREM2 potently inhibited mAβ42 self-polymerization in a dose-dependent manner (**Fig. 5.D**). These results demonstrate that TREM2 can inhibit Aβ42 aggregation and suggest that sTREM2 might perform this function *in vivo*.

**Fig. 5.**
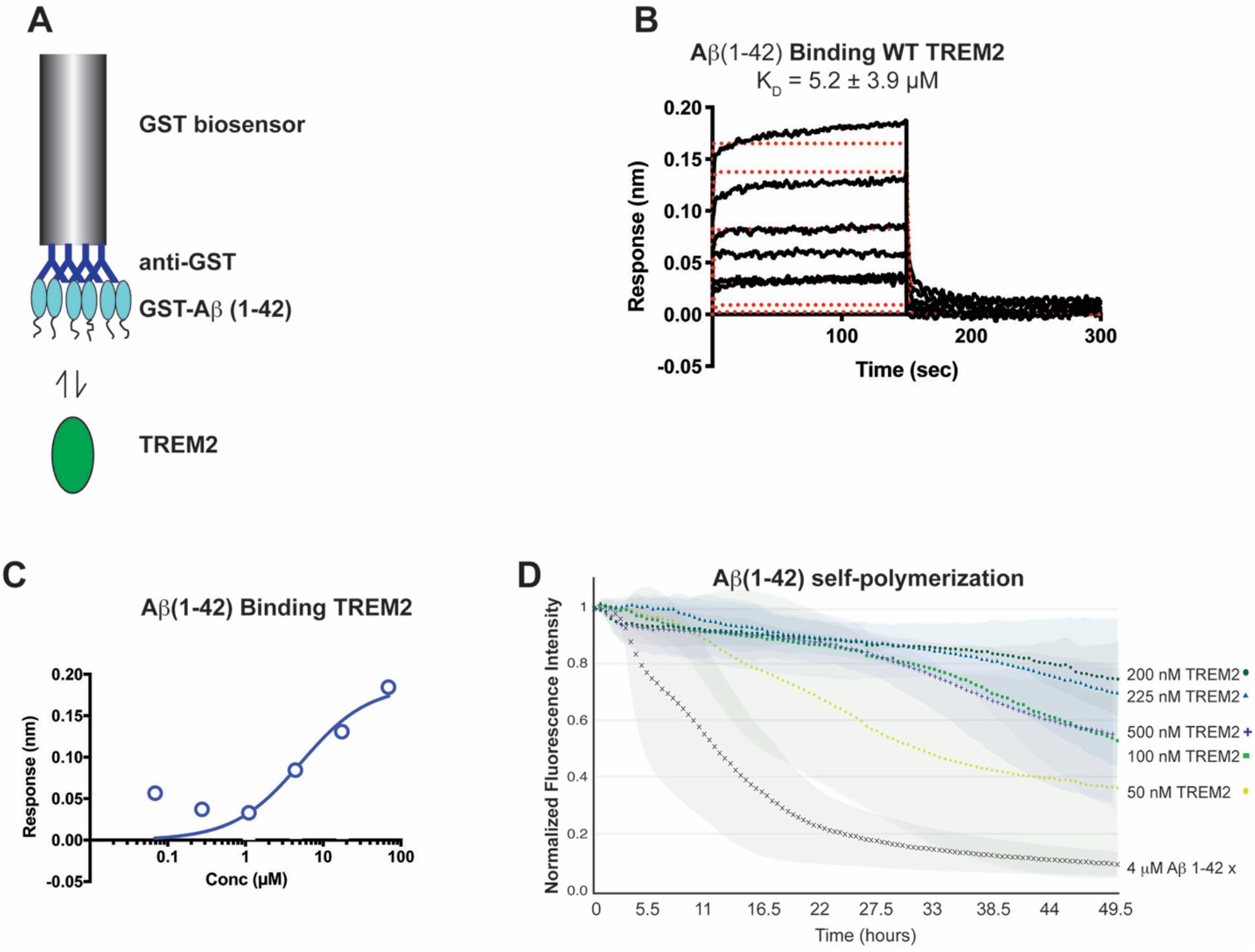
TREM2 binds monomeric Aβ42 and potently inhibits self-polymerization. **A)** Schematic for BLI experiment with GST-Aβ42 binding to purified TREM2. **B)** Binding response of TREM2 to GST-Aβ42. Data are double-reference subtracted using TREM2 binding to GST only. The concentration range for TREM2 was 0.068 - 70 μM. Affinity (K_D_) value determined by kinetic fits shown in red traces. **C)** Steady-state response of WT TREM2 binding to GST-Aβ42. Data are representative of three independent experiments. **D)** Self-polymerization of Aβ42 was monitored by fluorescence quenching of TMR-Aβ42. Markers represent the average of 3 runs while colored areas behind curves represent standard deviation.

## 3. Discussion

We conducted a rigorous molecular dissection of the TREM2-ApoE interaction and discovered a number of functionally relevant novel insights. First, our results show that TREM2 directly engages ApoE and does not require lipidation, demonstrating for the first time that the interaction is predominantly mediated by protein determinants (**Fig. 1**). Lipid loading of recombinant ApoE actually decreased the affinity (K_D_) of TREM2 interaction with ApoE3 from 440 nM to 1050 nM. ApoE particles with endogenous lipids derived from astrocytes also appeared to bind with similarly low affinity, although more comprehensive analysis is required. This decrease in affinity is likely due to either a decrease in avidity, since unloaded ApoE forms a concentration dependent monomer-dimer-tetramer mixture [34] while lipidated ApoE is a most likely a monomer [33], or because the large conformational change that occurs in ApoE upon engulfment of lipids [39, 40] alters the binding surface engaged by TREM2.

In terms of ApoE isoforms, we consistently found that TREM2 had the highest affinity for ApoE4, showing about a 1.5- and 2-fold higher affinity than for ApoE3 or ApoE2, respectively. In addition, in our experiments we found that the affinity of the TREM2-ApoE interaction (281 nM for TREM2-ApoE4) is considerably lower than what has been previously reported, whether it be by less quantitative estimates (6 - 9 nM) [22, 26] or by previous BLI studies (12 nM) [23]. For the latter study, the variance in these values likely arises from the different manners of TREM2 presentation in the experiments, as Fc-TREM2 fusion proteins were used. Fc-fusion proteins dimerize via the Fc portions, which increases avidity and usually contributes to biphasic binding, making kinetic analysis challenging. For our experimental design, we attempted to model as closely as possible the interactions as they would occur on the cell surface. Since TREM2 likely forms a 1:2 complex with DAP12 [41], it is likely presented as a monomer on the cell surface, so we immobilized monomeric TREM2 ectodomains in our BLI experiments. Thus, the differences in affinity reported are largely due to presentation of TREM2, with our values reported here more closely representing a single site-single site interaction.

We also showed that ApoE selectively binds TREM2 but does not appreciably interact with TREML2. GWASs have identified both TREM2 and TREML2 as modifiers for AD risk, albeit in opposite directions; point variants in TREM2 are correlated with increased AD risk [1, 2, 14], while variants in TREML2 are considered protective [28, 29]. It was recently shown that TREM2 and TREML2 play opposing roles in contributing to microglia proliferation and inflammatory response [30]. While the exact mechanisms for this are unknown, our results suggest that differential binding of ApoE could partially contribute to the observation of disparate functions for TREM2 compared to TREML2.

We also identified the major binding surface on TREM2 involved with engaging ApoE. We found that the AD-linked variants, located in the basic patch on TREM2, only moderately disrupt ApoE binding. Previous reports seem to support this finding, with a single-concentration BLI study finding a similar moderate loss of affinity to our observations [21] and another using titration BLI experiments to find almost no loss of affinity caused by these variants [23]. Our conclusion that the AD variants do not dramatically impact ApoE binding is strengthened by the discovery that a different surface on TREM2 is required for ApoE binding. We found that point mutations to a distal hydrophobic surface composed of the CDR1-3 loops on TREM2 reduced binding to ApoE by about 10-1000-fold. It is worth noting that a recent crystal structure of the TREM2 R47H variant [16] and results from molecular dynamics simulations of TREM2 R47H, R62H, and T96K variants [35] show that these AD risk variants induce structural instability in the CDR2 loop, which comprises a central portion of this hydrophobic patch. Thus, these structurally-induced impacts on the hydrophobic patch could explain why TREM2 basic patch mutations display moderate decreases in binding to ApoE. This hydrophobic patch also engages phospholipids in a recent co-crystal structure with TREM2 [16]. Future binding studies will need to validate that these hydrophobic site residues contribute to phospholipid binding and whether phospholipids and ApoE compete for binding to this site.

In addition, we have identified the major determinants on ApoE that mediate engagement of TREM2. The ApoE hinge region, in particular residues 192-238, most strongly contribute to ApoE binding. To our knowledge, this is the first time these residues have been implicated in direct protein-protein interactions. Our data suggest that the N-terminal domain of ApoE is involved in TREM2 binding as previously reported [42] but that the hinge region is the major determinant mediating interaction with TREM2 (**Fig. 4**). The importance of the major hinge region is consistent with our observation that lipidation alters binding to TREM2, as this region undergoes major conformational changes upon loading of lipids [43, 44]. These experiments extend the TREM2 epitope on ApoE and suggest that the TREM2-ApoE interaction could be selectively targeted. It should be noted that while the loss of avidity undoubtedly exaggerates the observed differences in binding between the full-length and ApoE3 1-238, it remains possible that the C-terminal domain contributes to direct interactions with TREM2. Future experiments with hydrogen-deuterium exchange mapping and structural studies of the TREM2-ApoE complex will be needed to clarify the full TREM2-ApoE binding surface.

Finally, we show that TREM2 directly binds mAβ42, and that this interaction inhibits Aβ42 oligomerization *in vitro*. This observation could indicate a potential protective *in vivo* function for sTREM2. Little is known regarding the function of sTREM2, but current observations suggest it plays a protective role in Alzheimer’s pathogenesis. For example, elevated levels of sTREM2 are associated with cognitive reserve in humans [10], and increasing sTREM2 in a mouse model of AD appears to ameliorate disease pathologies, including reducing amyloid plaque load and rescuing functional memory deficits [11]. Thus, understanding sTREM2 functions and how it is produced and regulated may shed new light on therapeutic targets [45]. Our data indicate that sTREM2 binds to mAβ42 and inhibits aggregation, which could be another route by which sTREM2 impedes AD development. Future structural and biophysical studies will need to identify the mAβ42 binding site on TREM2 so that sTREM2 variants lacking this function can be designed and utilized in *in vivo* experiments to address the potential role of this function in AD pathogenesis.

Altogether, our data along with previous published studies begin to clarify how TREM2 engages numerous molecules linked to AD-pathogenesis (**Fig. 6.A**). Our data show that TREM2 recognizes ligands using at least two separate surfaces and provide the first evidence for a role of the hydrophobic patch in recognizing a protein ligand (**Fig. 6.A,B)**. A basic patch of electropositive residues (including R47 and R62) is important for binding anionic ligands, such as cell-surface proteoglycans [15]. A separate hydrophobic surface is critical for ApoE binding, and also binds phospholipids (**Fig. 6.A**). Sequence analysis of TREM2 family proteins indicates that residues composing both the basic patch and the hydrophobic patch are conserved in human and mouse TREM2 but are not conserved in the overall TREM family, implying that these surfaces mediate TREM2-exclusive functions (**Fig. 6.C**). This is supported by our findings that TREML2, despite having a highly positive pI and (likely) a similar overall fold, does not engage ApoE. This identification of separate binding epitopes within the single TREM2 Ig domain will be crucial towards better understanding TREM2-ligand interactions, and will aid the rational design of therapeutics selectively targeting specific TREM2 functional surfaces. In addition, we show that TREM2 binds mAβ42 and inhibits polymerization. This observation suggests that sTREM2 could utilize this function to (at least partially) elicit its protective role in AD pathogenesis and highlights the need to understand regulation and production of sTREM2 by regulators such as MS4A4A [45].

**Fig. 6.**
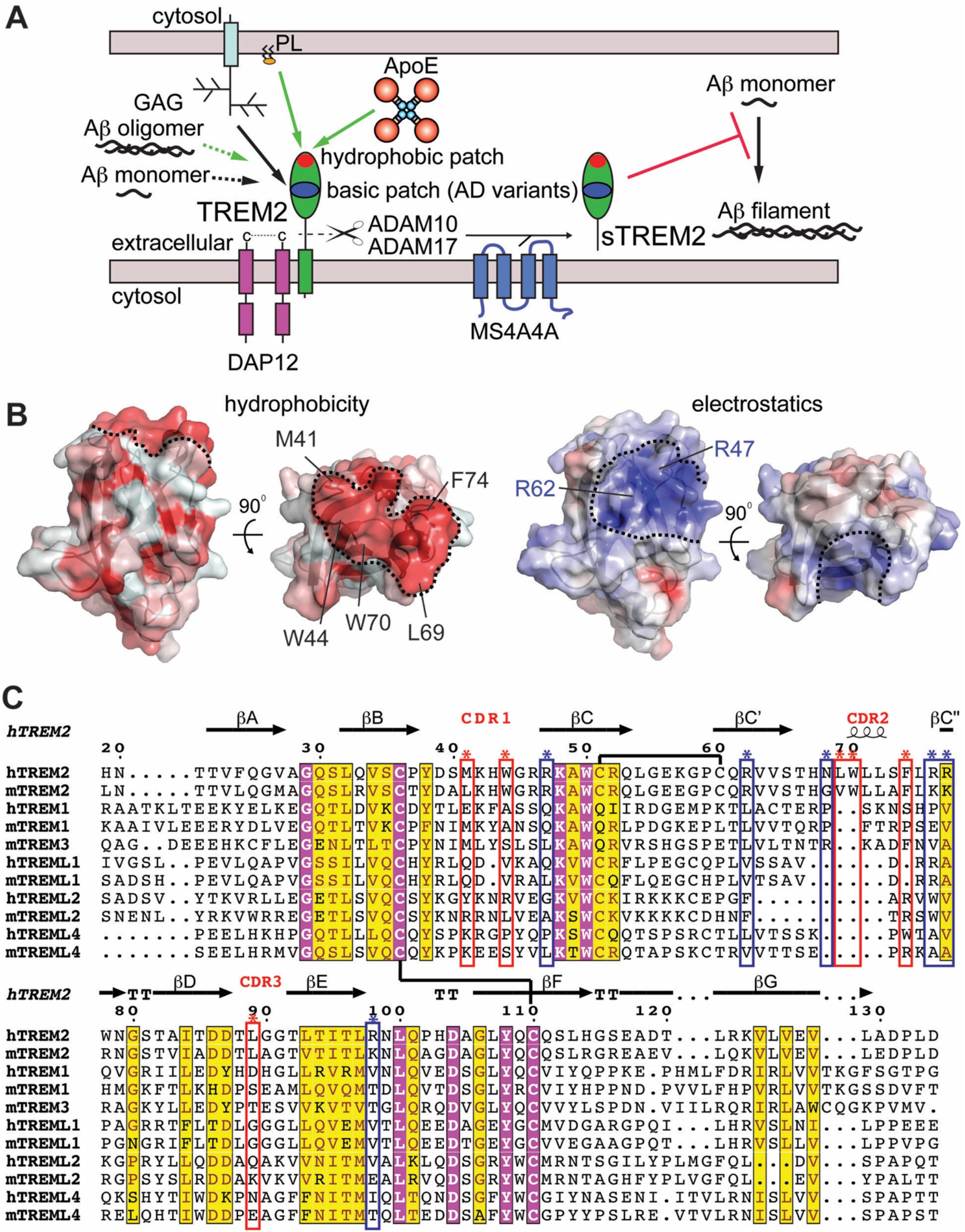
TREM2 has at least two unique ligand-binding surfaces. **A)** Graphic summary of known binding partners for TREM2 with proven or potential relevance to AD. Interaction surfaces mapped by either crystallography or mutations are shown as solid lines whereas direct interactions that have been shown but not mapped are shown as dashed lines. Green lines indicate interactions that result in TREM2 signaling through DAP12. Black lines indicate direct interactions that have not yet been shown to induce signaling. ApoE (this publication) and phospholipids [16] both engage the hydrophobic patch. Glycosaminoglycans (GAGs) engage the basic patch [15]. Soluble TREM2 (sTREM2) can be produced by proteolytic cleavage by ADAM10 and ADAM17 [6, 7] or by alternate transcripts [8]. Its levels appear to be modulated by MS4A4A [45]. This sTREM2 can engage monomeric Aβ42 and prevent polymerization. B) The TREM2 hydrophobic patch contrasted with the basic patch. Two orientations of the TREM2 (PDB 5ELI) protein surface are colored either by hydrophobicity (left) using the color_h pymol script where red is hydrophobic and white is hydrophilic, or by surface-accessible electrostatic potential (right) as in [15] where blue is basic and red is acidic. Scale is −6.0 kT/e (blue) to +6 kT/e (red). Key residues contributing to the hydrophobic surface are labeled on the top view. The hydrophobic and basic patches are outlined by the dashed black lines. C) Structure-based sequence alignment suggests the basic patch and hydrophobic patch are functional surfaces unique to TREM2 within the TREM family of human and mouse proteins. Chemically conserved and invariant residues are highlighted in yellow and magenta, respectively, using ESPript server. TREM2 secondary structure from PDB 5ELI [15] using the DSSP server. Disulfide bonds shown as black lines. TREM2 residues composing the hydrophobic patch are marked with red boxes and asterisks while the basic patch is denoted with blue boxes and asterisks.

## 4. Methods

### 4.1 Materials

BLI biosensors (streptavidin-coated (SA); and anti-GST (GST)) were purchased from Pall FortéBio. Black 96-well microplates were purchased from Griener Bio-One. Expi293 cells and Expi293 expression media were purchased from Thermo Fisher Scientific. Hype5 was purchased from Oz Biosciences. Dipalmitoylphosphatidylcholine (DPPC) was purchase from Avanti. Cholate was purchased from Sigma-Aldrich. Glutathione Sepharose 4B resin was purchased from GE Healthcare. Phosphatidylcholine assay kit was purchased from Sigma-Aldrich.

### 4.2 Production of recombinant ApoE

#### Expression and purification

Full-length ApoE2, E3 and E4 were expressed and purified generally as described previously [46]. Constructs were transformed into Rosetta2 (DE3) *E. coli,* and transformed cells were plated under antibiotic selection. Single colonies were selected, and starter cultures were grown overnight in Luria Broth (LB). The next day, 3-4 L of LB were grown at 37° C to an OD_600_ of ~ 0.8 before being induced with 0.5 mM IPTG for 5 hours at 30° C. Cells were pelleted and resuspended in 150 mM NaCl, 10 mM Tris pH 8.5, 50 mM K_2_HPO_4_, 5 mM imidazole, and 10 mM β-ME supplemented with lysozyme and DNAse. Cells were lysed by sonication and cleared by centrifugation at 16,000 x g. Supernatants were passed over NiNTA superflow resin (Qiagen). Resin was washed with 300 mM NaCl, 20 mM imidazole, and 50 mM K_2_HPO_4_ pH 8.0, and then eluted in 300 mM NaCl, 250 mM imidazole, and 50 mM K_2_HPO_4_, pH 8.0. The thioredoxin fusion was cleaved with PreScission Protease (a gift from Niraj Tolia) before being purified by size exclusion chromatography (SEC) in 20 mM HEPES pH 7.4 with 150 mM NaCl and 1 mM DTT. Truncation constructs were expressed and purified in a similar manner, except they were expressed without the thioredoxin fusion. Truncation constructs elute as apparent monomers by SEC, consistent with previous observations that the C-term domain mediates oligomerization [32].

#### Lipidation

ApoE3 was lipidated by complexing with DPPC similarly to that previously described [47]. DPPC:cholate lipoparticles were prepared by mixing cholate and DPPC at a ratio of 1.4:1 cholate:DPPC and incubating at 314 K for 1 hr with intermittent vortexing. SEC-purified ApoE3 was added at a ratio of 1:46 ApoE:DPPC for 10 min at RT followed by 1 hr at 314 K. Excess cholate was removed by buffer exchange through centrifugation and the protein was then re-purified on a Superose 6 Increase column (GE) in 20 mM HEPES pH 7.4, 150 mM NaCl, and 1 mM DTT at RT to remove ApoE aggregates along with excess lipid and cholate before BLI studies. A phosphatidylcholine assay kit (Sigma-Aldrich) confirmed ApoE lipidation.

#### Biotinylation

Purified full-length ApoE proteins were enzymatically biotinylated using BirA as described previously [15].

### 4.3 Expression and purification of GST-Aβ42

Aβ42 was expressed as a GST-fusion protein as previously described [48]. A codon-optimized sequence encoding the Aβ42 peptide was introduced into the pGEX-4T1 vector by Gibson block assembly using BamHI-XhoI restriction sites and was verified by sequencing. GST or GST-Aβ cultures were grown to an OD_600_ ~ 0.8, cold-shocked on ice for 20 min and then induced with 0.5 mM IPTG for 2 hr at 25 °C. Following induction, pelleted cells were resuspended in PBS pH 7.4 with 1 mM DTT, lysed by sonication and centrifuged at 16,000 x g for 30 min to pellet debris. The supernatant was collected and incubated with glutathione sepharose resin for 1 hour at 4° C. Beads were washed with PBS + 1 mM DTT and the proteins were eluted with 50 mM Tris pH 8.0, 10 mM reduced glutathione, and 1 mM DTT.

### 4.4 Expression, purification, and biotinylation of TREM proteins

TREM2 and TREML2 Ig domains or Ig domains with a BirA biotinylation tag were produced and biotinylated as previously described [15]. TREM2 Ig domain variants (M41D, M44D, L69D, W70D, and F74D) were produced by Gibson block assembly using AgeI/KpnI restriction digest sites in the pHLSEC vector and then transferred into the pHL-Avitag vector by restriction digestion and ligation [49]. All TREM2 constructs were verified by sequencing.

### 4.5 BLI experiments

BLI data were collected on an Octet RED384 system (FortéBio). Experiments were conducted in a running buffer of PBS supplemented with 0.1% BSA and 0.005% Tween-20. Assays with lipidated ApoE were conducted in the same buffer without Tween-20. Data were processed using double-reference subtraction. Steady-state affinities were determined by curve fitting the equilibrium responses in GraphPad Prism.

### 4.6 Prediction of TREM2-ApoE binding regions using computational methods

To identify regions on TREM2 that were likely to contain multiple protein-binding residues, the complete sequence of human TREM2 was run through the InteractionSites2 method [50] of PredictProtein [36]. InteractionSites2 is a neural network-based tool that identifies residues that are likely to be involved in protein-protein interactions. The predicted protein-binding residues that were located in the extracellular domain were then mapped onto the crystal structure of TREM2 to identify the functional regions of interest where the predicted residues clustered.

We then used an in-house hydropathy-matching program [51] to predict likely binding sites of the CDRs in TREM2 on ApoE. From the regions predicted by InteractionSites2 to contain multiple protein-binding residues (i.e. the Ig CDR1 and CDR2 loops of TREM2), two sequences, each identified as the 8 apical-most residues of a CDR loop, were selected. The sequences of the selected CDR loops were then converted into binary (+ or -) maps based on the sign of each amino acid’s hydrophobicity by the Kyte and Doolittle scale [52], and maps generated of each target region in both N→C and C→N orientations were scanned across a similar hydrophobicity map generated for whole sequence of ApoE. The percentage of complementary (+/-) pairs were recorded at each potential binding site on ApoE, the sites were then ranked, and those with >60% hydropathic complementarity to each of the CDR sequences of TREM2 were mapped onto the ApoE sequence to predict potential binding regions in ApoE for further experimental validation.

### 4.7 Aβ42 self-polymerization assays

Self-polymerization of tetramethyl-rhodamine labeled Aβ42 (TMR-Aβ42, WatsonBio Sciences) was quantified by fluorescence quenching as previously reported with minor modifications [38]. Assays were carried out in buffer consisting of 8 mM Na_2_HPO_4_, 2 mM KH_2_PO_4_ pH 7.4, 137 mM NaCl, 2.7 mM KCl, and 1 mM EDTA and were stirred at 25°C. Purified WT TREM2 was added at t = 0. Polymerization of TMR-Aβ42 was monitored using TMR fluorescence (λ_ex_ = 520 nm; λ_em_ = 600 nm).

## 5. Acknowledgements

This work was supported by the NIH [R01-HL119813] (T.J.B), [T32-GM007067] (D.L.K.), [T32-GM008361] (H.B.D), [T32-NS095775] (H.B.D), [K08-HL121168] (J.A.B), Knight Alzheimer’s Disease Research Center pilot grant [P50-AG005681-30.1] (T.J.B.), Alzheimer’s Association Research Grant [AARG-16-441560] (T.J.B.), the American Heart Association Predoctoral Fellowships [15PRE22110004 and 17PRE32780001] (D.L.K.), the Burroughs-Wellcome Fund Career Award for Medical Scientists (J.A.B.), and the Alzheimer’s Drug Discovery Foundation [GC-201804-2015209] (E.D.R. & Y.S.).

The authors declare no financial conflicts of interest.

## Author contributions

D.L.K., C.F., J.A.B., E.D.R., Y.S., and T.J.B. designed the study. D.L.K. and C.E.K. carried out BLI binding experiments and analysis under the supervision of J.A.B. and T.J.B. M.R.S. purified naturally lipid-loaded ApoE from astrocytes under the supervision of D.M.H. M.R.S. carried out BLI binding experiments using ApoE from astrocytes under the supervision of T.J.B. M.D.S.B. designed, carried out, and analyzed Aβ42 polymerization assays experiments under the supervision of C.F. D.F.S. carried out protein purification. S.S.N. developed expression constructs of TREM2 mutants. B.B. developed and provided ApoE expression constructs. H.B.D. carried out computational experiments under the supervision of E.D.R. and Y.S. D.L.K., M.D.S.B., C.E.K., H.B.D., M.R.S., D.M.H., C.F., J.A.B., E.D.R., Y.S., and T.J.B. wrote the manuscript.

## References

[1] Guerreiro R, Wojtas A, Bras J, Carrasquillo M, Rogaeva E, Majounie E, et al. TREM2 variants in Alzheimer’s disease. N Engl J Med. 2013;368:117–27.

[2] Jonsson T, Stefansson H, Steinberg S, Jonsdottir I, Jonsson PV, Snaedal J, et al. Variant of TREM2 associated with the risk of Alzheimer’s disease. N Engl J Med. 2013;368:107–16.

[3] Colonna M, Wang Y. TREM2 variants: new keys to decipher Alzheimer disease pathogenesis. Nat Rev Neurosci. 2016;17:201–7.

[4] Gratuze M, Leyns CEG, Holtzman DM. New insights into the role of TREM2 in Alzheimer’s disease. Mol Neurodegener. 2018;13:66.

[5] Ulland TK, Colonna M. TREM2 - a key player in microglial biology and Alzheimer disease. Nat Rev Neurol. 2018;14:667–75.

[6] Feuerbach D, Schindler P, Barske C, Joller S, Beng-Louka E, Worringer KA, et al. ADAM17 is the main sheddase for the generation of human triggering receptor expressed in myeloid cells (hTREM2) ectodomain and cleaves TREM2 after Histidine 157. Neurosci Lett. 2017;660:109–14.

[7] Thornton P, Sevalle J, Deery MJ, Fraser G, Zhou Y, Stahl S, et al. TREM2 shedding by cleavage at the H157-S158 bond is accelerated for the Alzheimer’s disease-associated H157Y variant. EMBO Mol Med. 2017;9:1366–78.

[8] Del-Aguila JL, Benitez BA, Li Z, Dube U, Mihindukulasuriya KA, Budde JP, et al. TREM2 brain transcript-specific studies in AD and TREM2 mutation carriers. Mol Neurodegener. 2019;14:18.

[9] Piccio L, Deming Y, Del-Aguila JL, Ghezzi L, Holtzman DM, Fagan AM, et al. Cerebrospinal fluid soluble TREM2 is higher in Alzheimer disease and associated with mutation status. Acta Neuropathol. 2016;131:925–33.

[10] Ewers M, Franzmeier N, Suarez-Calvet M, Morenas-Rodriguez E, Caballero MAA, Kleinberger G, et al. Increased soluble TREM2 in cerebrospinal fluid is associated with reduced cognitive and clinical decline in Alzheimer’s disease. Sci Transl Med. 2019;11.

[11] Zhong L, Xu Y, Zhuo R, Wang T, Wang K, Huang R, et al. Soluble TREM2 ameliorates pathological phenotypes by modulating microglial functions in an Alzheimer’s disease model. Nat Commun. 2019;10:1365.

[12] Zhong L, Chen XF, Wang T, Wang Z, Liao C, Wang Z, et al. Soluble TREM2 induces inflammatory responses and enhances microglial survival. J Exp Med. 2017;214:597–607.

[13] Wu K, Byers DE, Jin X, Agapov E, Alexander-Brett J, Patel AC, et al. TREM-2 promotes macrophage survival and lung disease after respiratory viral infection. J Exp Med. 2015;212:681–97.

[14] Jin SC, Benitez BA, Karch CM, Cooper B, Skorupa T, Carrell D, et al. Coding variants in TREM2 increase risk for Alzheimer’s disease. Hum Mol Genet. 2014;23:5838–46.

[15] Kober DL, Alexander-Brett JM, Karch CM, Cruchaga C, Colonna M, Holtzman MJ, et al. Neurodegenerative disease mutations in TREM2 reveal a functional surface and distinct loss-of-function mechanisms. Elife. 2016;5:e20391.

[16] Sudom A, Talreja S, Danao J, Bragg E, Kegel R, Min X, et al. Molecular basis for the loss-of-function effects of the Alzheimer’s disease-associated R47H variant of the immune receptor TREM2. J Biol Chem. 2018;293:12634–46.

[17] Kober DL, Brett TJ. TREM2-ligand interactions in health and disease. Journal of Molecular Biology. 2017.

[18] Wang Y, Cella M, Mallinson K, Ulrich JD, Young KL, Robinette ML, et al. TREM2 lipid sensing sustains the microglial response in an Alzheimer’s disease model. Cell. 2015;160:1061–71.

[19] Shirotani K, Hori Y, Yoshizaki R, Higuchi E, Colonna M, Saito T, et al. Aminophospholipids are signal-transducing TREM2 ligands on apoptotic cells. Sci Rep. 2019;9:7508.

[20] Song W, Hooli B, Mullin K, Jin SC, Cella M, Ulland TK, et al. Alzheimer’s disease-associated TREM2 variants exhibit either decreased or increased ligand-dependent activation. Alzheimers Dement. 2017;13:381–7.

[21] Yeh FL, Wang Y, Tom I, Gonzalez LC, Sheng M. TREM2 Binds to Apolipoproteins, Including APOE and CLU/APOJ, and Thereby Facilitates Uptake of Amyloid-Beta by Microglia. Neuron. 2016;91:328–40.

[22] Jendresen C, Arskog V, Daws MR, Nilsson LN. The Alzheimer’s disease risk factors apolipoprotein E and TREM2 are linked in a receptor signaling pathway. J Neuroinflammation. 2017;14:59.

[23] Lessard CB, Malnik SL, Zhou Y, Ladd TB, Cruz PE, Ran Y, et al. High-affinity interactions and signal transduction between Abeta oligomers and TREM2. EMBO Mol Med. 2018;10.

[24] Zhao Y, Wu X, Li X, Jiang LL, Gui X, Liu Y, et al. TREM2 Is a Receptor for beta-Amyloid that Mediates Microglial Function. Neuron. 2018;97:1023-31 e7.

[25] Zhong L, Wang Z, Wang D, Wang Z, Martens YA, Wu L, et al. Amyloid-beta modulates microglial responses by binding to the triggering receptor expressed on myeloid cells 2 (TREM2). Mol Neurodegener. 2018;13:15.

[26] Atagi Y, Liu CC, Painter MM, Chen XF, Verbeeck C, Zheng H, et al. Apolipoprotein E Is a Ligand for Triggering Receptor Expressed on Myeloid Cells 2 (TREM2). J Biol Chem. 2015;290:26043–50.

[27] Bailey CC, DeVaux LB, Farzan M. The Triggering Receptor Expressed on Myeloid Cells 2 Binds Apolipoprotein E. J Biol Chem. 2015;290:26033–42.

[28] Benitez BA, Jin SC, Guerreiro R, Graham R, Lord J, Harold D, et al. Missense variant in TREML2 protects against Alzheimer’s disease. Neurobiol Aging. 2014;35:1510 e19-26.

[29] Jiang T, Wan Y, Zhou JS, Tan MS, Huang Q, Zhu XC, et al. A Missense Variant in TREML2 Reduces Risk of Alzheimer’s Disease in a Han Chinese Population. Mol Neurobiol. 2017;54:977–82.

[30] Zheng H, Liu C-C, Atagi Y, Chen X-F, Jia L, Yang L, et al. Opposing roles of the triggering receptor expressed on myeloid cells 2 and triggering receptor expressed on myeloid cells-like transcript 2 in microglia activation. Neurobiology of Aging. 2016;42:132–41.

[31] de Freitas A, Banerjee S, Xie N, Cui H, Davis KI, Friggeri A, et al. Identification of TLT2 as an Engulfment Receptor for Apoptotic Cells. The Journal of Immunology. 2012;188:6381–8.

[32] Aggerbeck LP, Wetterau JR, Weisgraber KH, Wu CS, Lindgren FT. Human apolipoprotein E3 in aqueous solution. II. Properties of the amino- and carboxyl-terminal domains. Journal of Biological Chemistry. 1988;263:6249–58.

[33] Garai K, Baban B, Frieden C. Dissociation of apolipoprotein E oligomers to monomer is required for high-affinity binding to phospholipid vesicles. Biochemistry. 2011;50:2550–8.

[34] Garai K, Frieden C. The association-dissociation behavior of the ApoE proteins: kinetic and equilibrium studies. Biochemistry. 2010;49:9533–41.

[35] Dean HB, Roberson ED, Song YH. Neurodegenerative Disease-Associated Variants in TREM2 Destabilize the Apical Ligand-Binding Region of the Immunoglobulin Domain. Front Neurol. 2019;10.

[36] Yachdav G, Kloppmann E, Kajan L, Hecht M, Goldberg T, Hamp T, et al. PredictProtein--an open resource for online prediction of protein structural and functional features. Nucleic Acids Res. 2014;42:W337–W43.

[37] Raulin AC, Kraft L, Al-Hilaly YK, Xue WF, McGeehan JE, Atack JR, et al. The Molecular Basis for Apolipoprotein E4 as the Major Risk Factor for Late-Onset Alzheimer’s Disease. J Mol Biol. 2019;431:2248–65.

[38] Garai K, Verghese PB, Baban B, Holtzman DM, Frieden C. The binding of apolipoprotein E to oligomers and fibrils of amyloid-beta alters the kinetics of amyloid aggregation. Biochemistry. 2014;53:6323–31.

[39] Peters-Libeu CA, Newhouse Y, Hall SC, Witkowska HE, Weisgraber KH. Apolipoprotein E*dipalmitoylphosphatidylcholine particles are ellipsoidal in solution. J Lipid Res. 2007;48:1035–44.

[40] Peters-Libeu CA, Newhouse Y, Hatters DM, Weisgraber KH. Model of biologically active apolipoprotein E bound to dipalmitoylphosphatidylcholine. J Biol Chem. 2006;281:1073–9.

[41] Call ME, Wucherpfennig KW, Chou JJ. The structural basis for intramembrane assembly of an activating immunoreceptor complex. Nat Immunol. 2010;11:1023–9.

[42] Jendresen C, Årskog V, Daws MR, Nilsson LNG. The Alzheimer’s disease risk factors apolipoprotein E and TREM2 are linked in a receptor signaling pathway. Journal of Neuroinflammation. 2017;14:59.

[43] Chen J, Li Q, Wang J. Topology of human apolipoprotein E3 uniquely regulates its diverse biological functions. Proceedings of the National Academy of Sciences. 2011;108:14813–8.

[44] Hatters DM, Peters-Libeu CA, Weisgraber KH. Apolipoprotein E structure: insights into function. Trends Biochem Sci. 2006;31:445–54.

[45] Deming Y, Filipello F, Cignarella F, Cantoni C, Hsu S, Mikesell R, et al. The MS4A gene cluster is a key modulator of soluble TREM2 and Alzheimer’s disease risk. Sci Transl Med. 2019;11.

[46] Garai K, Mustafi SM, Baban B, Frieden C. Structural differences between apolipoprotein E3 and E4 as measured by (19)F NMR. Protein Sci. 2010;19:66–74.

[47] Newhouse Y, Peters-Libeu C, Weisgraber KH. Crystallization and preliminary X-ray diffraction analysis of apolipoprotein E-containing lipoprotein particles. Acta Crystallogr Sect F Struct Biol Cryst Commun. 2005;61:981–4.

[48] Zhang L, Yu H, Song C, Lin X, Chen B, Tan C, et al. Expression, purification, and characterization of recombinant human β-amyloid42 peptide in Escherichia coli. Protein Expression and Purification. 2009;64:55–62.

[49] Aricescu aR, Lu W, Jones EY. A time- and cost-efficient system for high-level protein production in mammalian cells. Acta crystallographica Section D, Biological crystallography. 2006;62:1243–50.

[50] Ofran Y, Rost B. ISIS: interaction sites identified from sequence. Bioinformatics. 2007;23:e13-e6.

[51] Fancy RM, Wang L, Schmid T, Zeng Q, Wang H, Zhou T, et al. Characterization of the Interactions between Calmodulin and Death Receptor 5 in Triple-negative and Estrogen Receptor-positive Breast Cancer Cells: AN INTEGRATED EXPERIMENTAL AND COMPUTATIONAL STUDY. J Biol Chem. 2016;291:12862–70.

[52] Kyte J, Doolittle RF. A simple method for displaying the hydropathic character of a protein. Journal of Molecular Biology. 1982;157:105–32.

[53] Chen J, Li Q, Wang J. Topology of human apolipoprotein E3 uniquely regulates its diverse biological functions. Proc Natl Acad Sci U S A. 2011;108:14813–8.

