## Supplemental Figures and Tables for "Functional insights from biophysical study of TREM2 interactions with ApoE and Aβ_1-42_"

***Supplemental methods***

***Cloning of ApoE constructs:***

Quickchange site directed mutagenesis introduced stop codons into the pET30(+) vector containing His-tagged monomeric ApoE (a gift from Jianjun Wang, described in [48]) at positions 170 and 239. ApoE3 1-191 was generated by PCR from the 1-238 template using the forward primer 5'-GGGAATTCCATATGCACCATCATCATCATCAT and reverse primer 5'-CCGCTCGAGTTACCGCACGCGGCCCTGTTCC and ligated into a pET23b vector with NdeI/XhoI restriction sites. Overlap extension PCR cloning was used to incorporate the BirA peptide and linker sequence (GLNDIFEAQKIEWHEGGSGGS) at the N-terminus of ApoE upon expression. Briefly, two initial PCR reactions created overlapping termini of linear insert containing BirA sequence at 5' end of APOE (N-terminal product) and a vector backbone. A second PCR reaction using the products from the initial PCR reactions created the final plasmid containing BirA-GGSGGS-ApoE4. Insert Primers: Sense - 5' CATTTTCGAG GCGCAAAAAA TTGAATGGCA TGAAGGAGGA AGCGGCGGTT CCAAGGTTGA ACAGGCTGTT GAAACTGAAC CGG 3', Antisense - 5' GCAAGCTTGT CGACGGAGCT CGAATTCAGT GATTGTCG 3' Vector Primers: Sense - 5' CGACAATCAC TGAATTCGAG CTCCGTCGAC AAGCTTGC 3', Antisense - 5' CCTCCTTCAT GCCATTCAAT TTTTTGCGCC TCGAAAATGT CATTCAGGCC GGGTCCTTGA AATAGCACTT CCAGGTCGTC G 3'. All constructs were sequence-verified.

**Supplemental figures**

Supplementary Figure 1

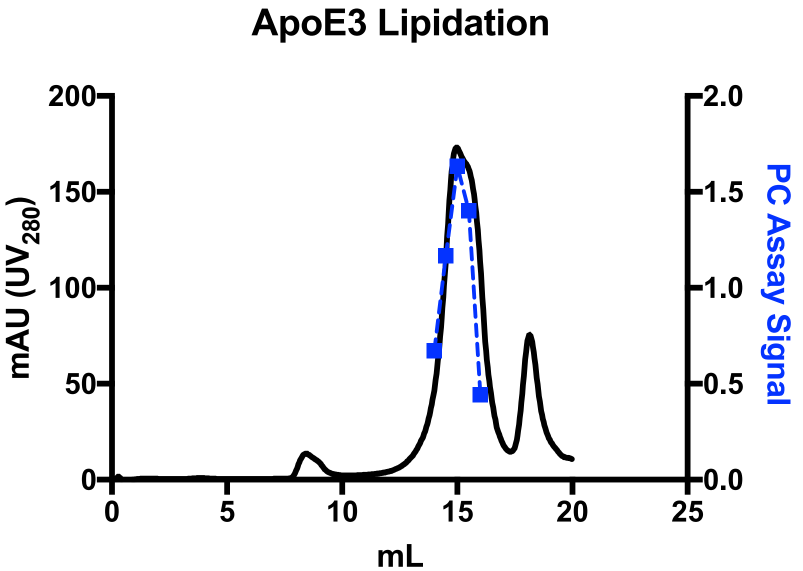


**Figure S1. SEC purification and analysis of lipidated ApoE3.** ApoE3 complexed with DPPC particles was purified by SEC (Superose 6) before BLI assays. The major peak was collected, and a PC assay signal (blue markers) from the collected fractions correlated with the UV peak, showing the presence of PC in the ApoE lipoparticles. Buffer alone and non-lipidated ApoE3 had no signal (not shown).

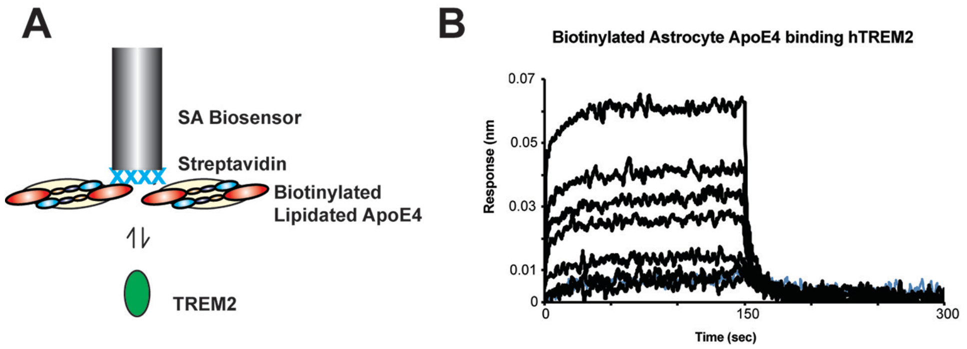


**Figure S2. Biotinylated astrocyte derived ApoE4 binding to TREM2 using BLI.** Lipid-loaded ApoE4 affinity purified from astrocytes was biotinylated using EZ-link PEG4 biotin and immobilized on a streptavidin biosensor. Purified TREM2 was in the wells at a concentration range of 3-200 μM. Sensorgrams were subtracted by double referencing. Lipid composition of ApoE lipoparticles did not appear to dramatically enhance binding to TREM2, as astrocyte-derived ApoE4 lipoparticles bound similarly to ApoE4:DPPC particles. However, the analysis was not exhaustive and may have been influenced by the orientation of the experiment.

Supplementary Figure 3.

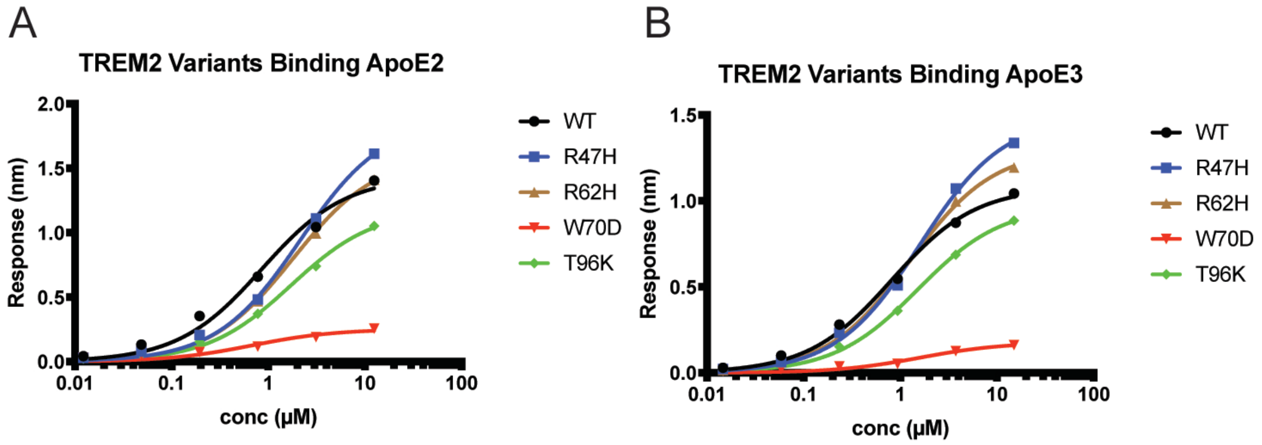


**Figure S3.** Steady-state binding of **A)** ApoE2 and **B)** ApoE3 to WT and TREM2 variants immobilized on SA BLI biosensors.

Supplementary Figure 4

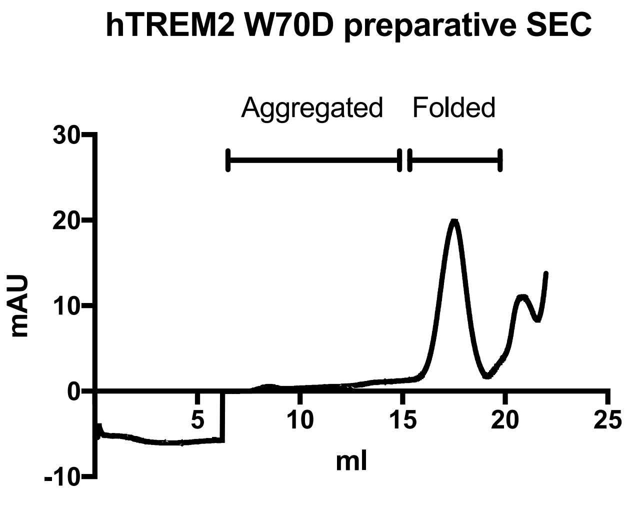


**Figure S4. W70D mutations does not cause aggregation as assessed by SEC.** W70D preparative SEC trace. Variant elutes in the fractions corresponding to folded TREM2, with minimal aggregated protein.

Supplementary Figure 5

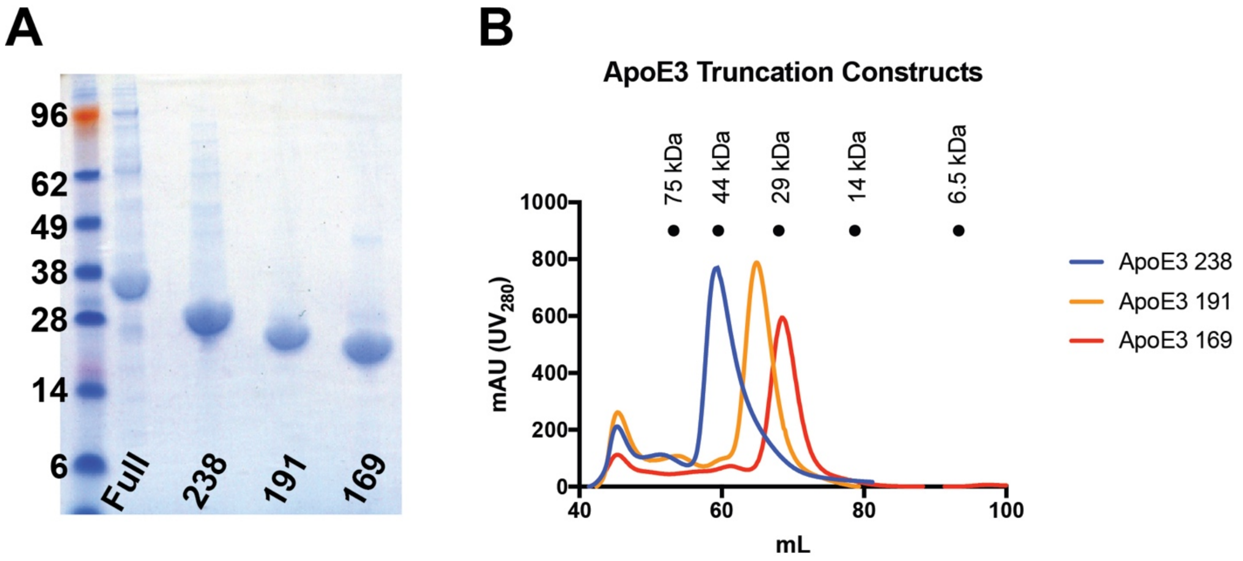


C

| ApoE Construct | MW (calc) | MW (SEC) | Relative to calculated MW |
|---|---|---|---|
| 238 | 28700 | 49147 | 1.71 |
| 191 | 23500 | 34121 | 1.45 |
| 169 | 20900 | 27578 | 1.32 |

**Figure S5. Purification and behavior of ApoE3 truncations. A)** Purity of Full-length ApoE3 truncations used in Figure 3 is shown by SDS PAGE. **B)** SEC traces of ApoE3 truncations and elution volumes of molecular standards. **C)** Table of calculated and experimental (SEC) MW of ApoE truncations. All truncations run about 1.5x their expected size, below a dimer size. The apparent increase is likely due to their elongated, non-globular structure.

Supplementary Figure 6

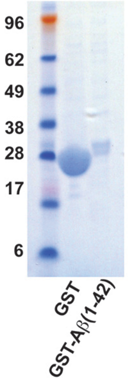


**Figure S6.** SDS-PAGE gel of GST and GST-Aβ used in the experiment shown in Figure 5C,D. Molecular weight markers are labeled in kDa.

Supplementary Table 1

| Region (Residues) | | CDR Forward | | | CDR Reverse | | |
|---|---|---|---|---|---|---|---|
| TREM2 | ApoE | Match | Start | End | Match | Start | End |
| CDR1 (39-46) | LDLR Region (130–149) | **75.00%** | **134** | **141** | **62.50%** | **134** | **141** |
| | Hinge Region (192–238) | **62.50%** | **192** | **199** | **75.00%** | **192** | **199** |
| CDR2 (69-76) | LDLR Region (130–149) | 87.50% | 134 | 141 | 87.50% | 135 | 142 |
| | | **87.50%** | **137** | **144** | **100.00%** | **138** | **145** |
| | | 87.50% | 140 | 147 | 75.00% | 141 | 148 |
| | Hinge Region (192–238) | | | | 75.00% | 192 | 199 |
| | | | | | 62.50% | 193 | 200 |
| | | 75.00% | 194 | 201 | 75.00% | 195 | 202 |
| | | 62.50% | 199 | 206 | 62.50% | 199 | 206 |
| | | 62.50% | 200 | 207 | 62.50% | 200 | 207 |
| | | | | | 62.50% | 201 | 208 |
| | | 75.00% | 202 | 209 | 75.00% | 203 | 210 |
| | | | | | 62.50% | 204 | 211 |
| | | 62.50% | 206 | 213 | 62.50% | 206 | 213 |
| | | 62.50% | 208 | 215 | 62.50% | 208 | 215 |
| | | 62.50% | 210 | 217 | 62.50% | 210 | 217 |
| | | | | | 75.00% | 215 | 222 |
| | | **87.50%** | **217** | **224** | **87.50%** | **218** | **225** |
| | | 75.00% | 220 | 227 | 75.00% | 221 | 228 |
| | | 75.00% | 222 | 229 | 75.00% | 223 | 230 |
| | | **87.50%** | **225** | **232** | **87.50%** | **226** | **233** |
| | | | | | 75.00% | 229 | 236 |

Rows with colored text represent top the predicted binding site(s) between a pair of regions. Colors match those used in Figure 2.

**Table S1.** Binding sites on the LDLR-binding region and hinge region of ApoE predicted by hydropathy mapping to be >60% complementary to forward or reverse peptides of the TREM2 hydrophobic surface.
